# A complex tissue-specific interplay between the Arabidopsis transcription factors AtMYB68, AtHB23, and AtPHL1 modulates primary and lateral root development and adaptation to salinity

**DOI:** 10.1101/2022.12.18.520920

**Authors:** Fiorella Paola Spies, María Florencia Perotti, Yuhan Cho, Chang Ig Jo, Jong Chan Hong, Raquel Lía Chan

## Abstract

- Adaptation to soil is a well-regulated process vital for plant life. AtHB23 is a homeodomain-leucine zipper I transcription factor (TF), previously revealed as crucial for plant survival in front of salinity conditions. We wondered whether this TF has partners to achieve this essential function.
- A TF cDNA library screening, Y2H, BiFC, and CoIP assays were complemented with expression analyses and phenotypic characterizations of silenced, mutant, overexpressor, and crossed plants in normal and salinity conditions.
- We revealed that AtHB23, AtPHL1, and AtMYB68 interact with each other, modulating root development and salinity response. The encoding genes coexpress in specific root tissues and developmental stages. In normal conditions, *amiR68* silenced plants have less initiated roots, the opposite phenotype to that showed by *amiR23* ones. AtMYB68 and AtPHL1 play contrary roles in lateral root elongation. Under salinity, where AtHB23 plays a crucial positive function, AtMYB68 cooperates with it, whereas AtPHL1 obstructs its action impacting survival ability and supporting the complex interaction between AtHB23, AtPHL1, and AtMYB68 in the primary and lateral roots. The root adaptation capability was associated with the amyloplast state.
- We identified new molecular players that through a complex relationship determine Arabidopsis root architecture and survival ability in salinity conditions.

## INTRODUCTION

Plants’ adaptation to the environment is a finely regulated process involving different biomolecules and levels of modulation. Roots are the anchorage organs that firstly sense changes in the soil and accordingly accelerate or arrest their primary or lateral growth and development for better adaptation, optimizing water and nutrient uptake (de Dorlodot *et al*., 2007; Waidmann *et al*., 2020). They enable plant adaptation to unfavorable environments by integrating different cues and balancing growth and development (Schachtman and Goodger, 2008).

Phytohormones like ABA and auxin play crucial roles in such an adaptation. Under salinity stress, the growth of the primary root is inhibited concomitantly with a decrease in the auxin content in the tip. Auxin transport toward roots is carried out in the central cylinder by AUX1, LAX1, LAX2, and LAX3 carriers exhibiting specific expression patterns impacting the phytohormone content in each tissue (Friml *et al*., 2003; Péret *et al*., 2012; Swarup and Péret, 2012). In the tip, NaCl regulates AUX1 and PIN2 (Liu *et al*., 2015). Low NaCl concentrations (≤50 mM) generate lateral root (LR) growth, whereas higher ones repress it, and these changes depend on auxin transport and distribution (Zolla *et al*., 2010).

Salinity stress and auxin content and distribution are closely related. Starch granules synthesis happens in the columella cells and is a process affected by salinity, affecting the gravitational potential energy (Leitz *et al*., 2009; Korver *et al*., 2020; Zhang *et al*., 2019).

Molecular events modulating root architecture and plasticity in changing and harmful conditions involve the participation of transcription factors (TF). These regulatory proteins play clue roles, usually activating or repressing entire transcriptional programs. Plants’ TF are abundant (about 1600 in Arabidopsis and 1500 in rice) and classified in families and subfamilies, mainly according to their DNA-binding domains, which determine their target specificity (González, 2016; Hong, 2016). For example, in LR development, proteins from the ARF (Auxin Response Factors) and LBD (Lateral Organ Boundaries (LOB) Domain) families have been assigned as master regulators (Friml *et al*., 2003; Xu *et al*., 2020; Banda *et al*., 2020; Lavenus *et al*., 2013; Lee *et al*., 2009).

The homeodomain (HD) TF family is large; its members differ in size, gene structure, localization of the HD, and other features (Capella *et al*., 2016). This fact generated a further classification of subfamilies. Among them, the HD-Zip subfamily is composed of proteins having a leucine zipper (LZ) downstream and adjacent to the HD and divided into four groups named I to IV. Functional divergence between members of subfamily I, and neofunctionalization, was explained by different uncharacterized motifs in the amino- and carboxy-termini of such proteins (Arce *et al*., 2011; Capella *et al*., 2014). Among these motifs, the AHA, at the 3’ end of the carboxy terminus, interacts with the basal transcriptional machinery (Capella *et al*., 2014). However, the deletion of the AHA motif to avoid transactivation in yeast-two-hybrid (Y2H) assays indicated that the proteins still interact with others. Such was the case of AtHB23 (AtHB23ΔAHA), used as bait to screen an Arabidopsis TF library, which revealed the interaction with four different proteins (Spies *et al*., 2022). They were three belonging to the large MYB family, AtMYB68 (At5g65890), AtPHL1 (At5g29000), and AtMYB12 (At2g47460), and AtWRKY43 (At2g46130; Spies *et al*., 2022).

AtHB23 is a member of group α (Henriksson *et al*., 2005) or clade V (Arce *et al*., 2011). This gene was expressed in the lateral root primordium, acting as a negative regulator of LR initiation and in the tip of primary roots involved in the salinity response (Perotti *et al*., 2019). It was directly regulated by ARF7/19, and *LAX3* and *LBD16* are its targets (Perotti *et al*., 2019, 2022).

*AtMYB68* was expressed during LR development and transcriptionally induced by high temperatures (Feng *et al*., 2004). However, its role in LR development remains uncertain. It was involved in several regulatory networks controlling the development, metabolism, and responses to biotic and abiotic stresses (Dubos *et al*., 2010). A few members of this family, such as AtMYB52, 53, 56, and 87, have been linked to LR development. Moreover, they participate in the intricate auxin-responsive network of TF (Lavenus *et al*., 2015).

AtPHL1 belongs to the MYB-CC subfamily having 15 members characterized by the MYB domain and a coiled-coiled (CC) domain (Rubio *et al*., 2001). Although AtPHL1 was thought redundant with AtPHR1, the most studied protein of this family, such redundancy was only partial and related to Pi starvation. PHR1 was identified as the master modulator of Pi-deficiency responses, inducing transcription of Pi-starvation genes, whereas the aberrant expression of *phl1* was only mildly affected, indicating a minor role in such an event (Bustos *et al*., 2010). The analysis of the double mutant *phr1/phl1* revealed that both genes participated in iron homeostasis regulation (Bournier *et al*., 2013). Besides their participation as players in the Pi-starvation response in several plant species, the role of MYB-CC in root development or stress responses remains largely unknown.

It was previously reported that *AtPHL1* and *AtHB23* are coexpressed in the pedicel-silique nodes and the funiculus, interacting to promote sucrose transport (Spies *et al*., 2022). In this work, we investigated the interplay between AtHB23, AtMYB68, and AtPHL1 in root development. We found that AtHB23 interacts with both MYB TFs, in yeast and plants. Moreover, these partners interacted between them too. The three TF coexpressed in specific cell groups and developmental stages of the primary and lateral roots. Under control and salinity conditions, they play cooperative and opposite roles depending on the situation.

## MATERIALS AND METHODS

### Plant material and growth conditions

*Arabidopsis thaliana* plants (Col-0 ecotype) were grown on Klasmann Substrat N° 1 compost (Klasmann-Deilmann GmbH, Germany) in a growth chamber at 22-24 °C under long-day (16/8 h light/dark cycles) conditions, with a light intensity of approximately 120 μmol m^−2^ s^−1^ in 8 × 7 cm pots. Four plants were planted per pot unless stated differently.

Transgenic plants carrying *AUX/LAX* promoters fused to *GUS*, previously described (*prAUX1:GUS*: Marchant *et al*. 1999, 2002; *prLAX1:GUS*: Bainbridge *et al*., 2008, and *prLAX3:GUS*: Swarup *et al*. 2008), were generously gifted by Dr. Swarup’ lab.

Arabidopsis mutant lines *phl1*-1 and −2 (SAIL_731_B09 and (SALK_079505.20.10) and *DR5:GFP* transgenic plants were obtained from the Arabidopsis Biological Resource Center (ABRC) stock. *AtHB23*-silenced plants (*amiR23*), *prAtPHL1:GUS, prAtHB23_L_:GUS, and prAtHB23_S_:GUS* were previously described (Perotti *et al*., 2019; Spies *et al*., 2022; Perotti *et al*., 2022).

### Genetic constructs used for plant transformation

#### 35S:AtMYB68

the vector *pENTER223* containing the coding sequence of *AtMYB68* (G22683) was purchased and recombined in the *pFK247* plasmid using the Gateway^®^ (Invitrogen) system.

#### 35S:AtMYB68:GFP

using as probe the construct *pENTER223-AtMYB68* (see above), the coding sequence corresponding to AtMYB68 was amplified with specific oligonucleotides (Supplementary Table S1). The amplification product was introduced in the *pGEM T-easy* vector and then subcloned in the plasmid *pENTER3C* between the *BamHI* and *XbaI* sites. By Gateway^®^ (Invitrogen) recombination, it was finally cloned in the destiny vector *pFK248*.

#### 35S:amiR68

the design of amiRMYB68 was carried out with the WMD3 software (Web MicroRNA Designer; wmd3.weigelworld.org) (Schwab *et al*., 2006). Four oligonucleotides (Supplementary Table S1), the endogenous mi319a, and the binary vector *pNB47* were used in an overlapping PCR. The amplification product was cloned in the *pGEM T-easy* vector and then in the *pENTER3C* previously restricted with *BamHI* and *EcoRI*. Finally, the amiRMYB68 was introduced in the *pKGWFS7* destiny plasmid using the Gateway^®^ (Invitrogen) recombination system,

#### PrMYB68:GUS:GFP

A 2377 bp fragment upstream of the ATG and corresponding to the promoter region of AtMYB68 was amplified by PCR using genomic DNA and specific oligonucleotides (Supplementary Table S1). The PCR product was cloned into the *pGEM-Teasy* vector, restricted with *BamHI* and *XhoI*, and finally recombined into the *pKGWFS7* destiny plasmid using the Gateway^®^ (Invitrogen) system.

### Genetic constructs used for BiFC and colocalization analyses

To perform PCR amplifications, cDNA was used as a probe, and specific oligonucleotides were listed in Supplementary Table S1.

#### AtHB23:GFP, Myc-AtHB23, Venus-N:AtHB23, and Venus-C:AtHB23

A 765 bp fragment starting from the ATG (TSS) of the *AtHB23* gene was amplified with specific oligonucleotides (Supplementary Table S1) and cloned into the pMDC83 (Ka+) Gateway^®^ vector system (Invitrogen), the Myc-pBA vector bearing the amino-terminal six-myc tag, the pDEST-GWVYNE vector, and the pDEST-GWVYCE vector (GATEWAY-BiFC vectors; Gehl *et al*., 2009), respectively.

#### MYB68:RFP, HA-AtMYB68, Myc-AtMYB68, Venus-N:AtMYB68, and Venus-C:AtMYB68

A 1122 bp fragment starting from the ATG (TSS) of the *AtMYB68* gene was amplified by PCR using cDNA and specific primers (Supplementary Table S1). The amplified PCR product was cloned into the pH7RWG2 (Spec+) Gateway^®^ vector system (VIB-UGent Center for Plant Systems Biology), the HA-pBA vector bearing in the amino-terminal a tri-haemagglutinin tag, the HA–pBA vector, the pDEST-^GW^VYNE, and pDEST-^GW^VYCE vectors, respectively.

#### HA-AtPHL1, Venus-C:AtPHL1, and Venus-N:PHL1

A 1239 bp fragment starting from the ATG (TSS) of the gene *AtPHL1* was amplified by PCR and cloned into the HA–pBA vector, the pDEST-^GW^VYCE, and pDEST-^GW^VYNE (Ka^+^) vector, respectively.

#### NLS-RFP

An *NLS* signal sequence tagged *NLS-RFP* was obtained from Prof. Sang Yeol Lee’s lab (Gyeongsang National University, Jinju).

#### Venus-N:AtHY5

A 540 bp fragment starting from ATG (TSS) of the gene AtHY5 was amplified and cloned into pDEST-^GW^VYNE vector.

#### Venus-C:AtSTO

A 744 bp fragment starting from the ATG (TSS) of the gene AtSTO was amplified by PCR and cloned into the pDEST-vector.

#### AD:PHL1, BD:PHL1, AD:AtHB23ΔAHA and BD:AtHB23ΔAHA

used for yeast two-hybrid assays were previously described (Spies *et al*., 2022).

### Arabidopsis stable transformation

Stable transformations of Arabidopsis plants were performed via a floral dip procedure as previously described (Clough and Bent, 1998). *Agrobacterium tumefaciens* strain, LBA4404, carrying the constructs described below, was used to transform. The selection was performed on the basis of their resistance with the appropriate selector chemical (Basta 50 mg/l or kanamycin 50 mg/l).

Transgene insertions were verified by PCR using genomic DNA as a template and specific oligonucleotides (Supplementary Table S1). Three/four positive independent lines were further reproduced and homozygous T3 and T4 plants were used to further analyses.

### Plant crosses

Mutant plants *phl1-1, phl1-2, amiR68-1*, and *amiR68-5* were fertilized with pollen from *prAUX1:GUS, prLAX1:GUS*, and *DR5:GUS* genotypes, and then selected by the corresponding selector chemical, depending on the donor (kanamycin resistance for the constructs bearing the promoters of *LAX1* and *AUX1*).

*AmiR23* and AT23 plants were fertilized with pollen from *phl1-1, phl1-2, OEPHL1-14, OEPHL1-16, amiR68-1*, and *amiR68-5* genotypes, *whereas* AT68-5 plants with that from *amiR68-1* and *amiR68-5* genotypes.

### Yeast two-hybrid screening

A truncated version of AtHB23 (AtHB23ΔAHA, Capella *et al*., 2014) was used as bait for yeast two-hybrid screening as previously described (Spies *et al*., 2022).

### *Nicotiana benthamiana* transient transformation for colocalization and BiFC analyses

*Nicotiana benthamiana* leaves were transformed by infiltration with a syringe containing 10 mM MES, 0.1 mM acetosyringone, 10 mM MgCl_2_, and cultured *A. tumefaciens* GV3101 (DO_600_: 0.3) previously transformed with the constructs indicated in the corresponding figures and mixed with *A. tumefaciens* cells transformed with the silencing inhibitor p19. Two days after infiltration, samples were harvested starting 2 h before the end of the day and used for visualization under confocal microscopy (FLUOVIEW FV3000-Olympus confocal laser microscope). Samples were excited using a 514 nm laser, and emission was detected in two channels: 520-530 nm for Venus and 540-600 nm for lignin autofluorescence.

### Co-immunoprecipitation

*Nicotiana benthamiana* leaves were infiltrated with Agrobacteria GV3101, at a final OD_600_nm = 0.5. Proteins were extracted using a buffer composed by 100 mM Tris-HCl (pH 7.4), 150 mM NaCl, 1mM EDTA (pH 8.0), 0.1% NP40, protease inhibitor (5 μg/mL Leupeptin, 1 μg/mL Aprotinin, 5 μg/mL Antipain, 1 μg/mL Pepstatin A, 5 μg/mL Chymostatin, 3 mM DTT, 100 μM PMSF, 1.5 mM Na_3_VO_4_, 2 mM NaF, 50 μM MG132), and coimmuniprecipitated by Dynabeads™ Protein A Immunoprecipitation Kit (Invitrogen™, Catalog No.10006D) with Myc-Tag (9B11) Mouse mAb (Cell Signaling Technology^®^, Catalog No. 2276S). Proteins were separated with 10% SDS-PAGE, then transferred to Immobilon^®^-P PVDF Membrane (Merck Millipore Ltd., Catalog N° IPVH00010) using a Trans-Blot^®^ Turbo™ Transfer System (Bio-Rad, Catalog No. #1704150). The immunoblots were detected by anti-HA-HRP (Roche, Code:12013819001) and anti-Myc-Tag (9B11) Mouse mAb (HRP Conjugate; Cell Signaling Technology^®^, Catalog No. 2040S) after 2 h incubation in room temperature. Images were taken by ChemiDoc™ MP Imaging System (Bio-Rad, Catalog N° #12003154).

### Root phenotyping

For root phenotyping: seeds (Col 0, mutant, and overexpressor plants) were surface sterilized and placed at 1 cm from the top of square Petri dishes (12 × 12 cm) for 3 days at 4 °C before placing the dishes in the growth chamber at 22-24 °C under long-day (16/8 light/dark cycles) with a light intensity of approximately 110-120 μmol m^−2^ s^−1^. The growing medium was Murashige-Skoog supplemented with vitamins (MS, PhytoTechnology Laboratories™).

For roots surveys, photographs series were taken and analyzed with ImageJ and RootNav free softwares.

### GUS histochemistry

*In situ* assays of GUS activity were performed essentially as described by Jefferson *et al*. (1987) with little modifications (Ribone *et al*., 2015).

### RNA isolation and expression analyses by RT-qPCR

Total RNA for real-time RT-PCR was isolated from rosette leaves of 25-day-old plants or 5-8 day-old roots using Trizol^®^ reagent (Invitrogen, Carlsbad, CA, USA) according to the manufacturer’s instructions. Total RNA (1 μg) was reverse-transcribed using oligo(dT)_18_ and M-MLV reverse transcriptase II (Promega, Fitchburg, WI, USA).

Quantitative real-time PCR (qPCR) was performed using an Mx3000P Multiplex qPCR system (Stratagene, La Jolla, CA, USA) as described before (Mora *et al*., 2022) and using the primers listed in Table S1. Transcript levels were normalized by applying the ΔΔCt method. Actin transcripts (*ACTIN2* and *ACTIN8*) were used as internal standards. Three biological replicates, obtained from 3 individual plants and tested by duplicate, were used to calculate the standard deviation.

### Amyloplasts Staining and Light Microscopy Observation

To observe the amyloplasts in the columella cells of the root tips, 15-20 Arabidopsis roots (5-day-old) were dipped in Lugol staining solution (Sigma-Aldrich) for 8-10 min, washed with distilled water, and then observed in an Eclipse E200 Microscope (Nikon, Tokyo, Japan, https://www.nikon.com/) equipped with a Nikon Coolpix L810 camera. For salinity treatments, seedlings were transferred to Petri dishes with MS supplemented with 150 mM NaCl for seven-eight additional hours, according to the method described by Sun *et al*., (2008).

### Fluorescence Microscopy

For confocal imaging, roots from different genotypes were treated with 10 μg/ml propidium iodide, rinsed with a drop of distilled water, and examined and imaged using a confocal inverted microscope (Confocal LEICA TCS SP8). The observations were done using a 20 × objective, a 514 nm excitation line laser for propidium iodide (18% intensity), and appropriate emission at 498 nm – 532 nm, bandpass filters.

### Statistical analysis

The evaluation of phenotypic characteristics such as primary root length, total lateral root length, number of initiated and emerged secondary roots, and number of cells, as well as quantitative PCR, was statistically analyzed using one-way analysis of variance (ANOVA) considering genotype as the main factor.

Significant differences between means were analyzed using Tukey’s posthoc comparison and indicated by different letters. The number of biological replicates for each assessment is indicated in the corresponding figures.

### Accession codes

AT1G26960 (AtHB23); AT5G65790 (AtMYB68), AT5gG29000 (AtPHL1), AT2G38120 (AUX1), AT5G48300 (ADG1), AT5G51820 (PGM), AT1G10760 (GWD), AT3G23920 (BAM1).

## RESULTS

### The interaction between transcription factors AtHB23, AtPHL1, and AtMYB68 was validated in yeast, *in vitro*, and *in planta*

Homeodomain-leucine zipper transcription factors exhibit uncharacterized motifs in their amino and carboxy termini, potentially interacting with other specific proteins (Arce *et al*., 2011). Such was the case of AtHB23, used as bait to screen an Arabidopsis TF library, allowing identifying four putative partners. In previous work, we corroborated the interaction between AtHB23 and AtPHR1-like1 (thereafter, AtPHL1) by carrying out independent Y2H and BiFC (bimolecular fluorescence complementation) assays (Spies *et al*., 2022). Moreover, the functional meaning of such an interaction was studied in conductive tissues and siliques (Spies *et al*., 2022), whereas that with AtMYB68 (At5g65890) remained unexplored. Hence, we decided to continue the study with this AtHB23 putative partner.

Firstly, we verified the interaction between AtHB23 and AtMYB68 by independent Y2H and β-galactosidase activity assays (Figure 1a). To further validate the relationship *in planta*, we examined their subcellular localization and determined that they co-localized in the nucleus (Figure 1b). Moreover, their interaction was confirmed by BiFC assays in *Nicotiana benthamiana* leaves (Figure 1c and Supplementary Figure S1).

**Figure 1.**
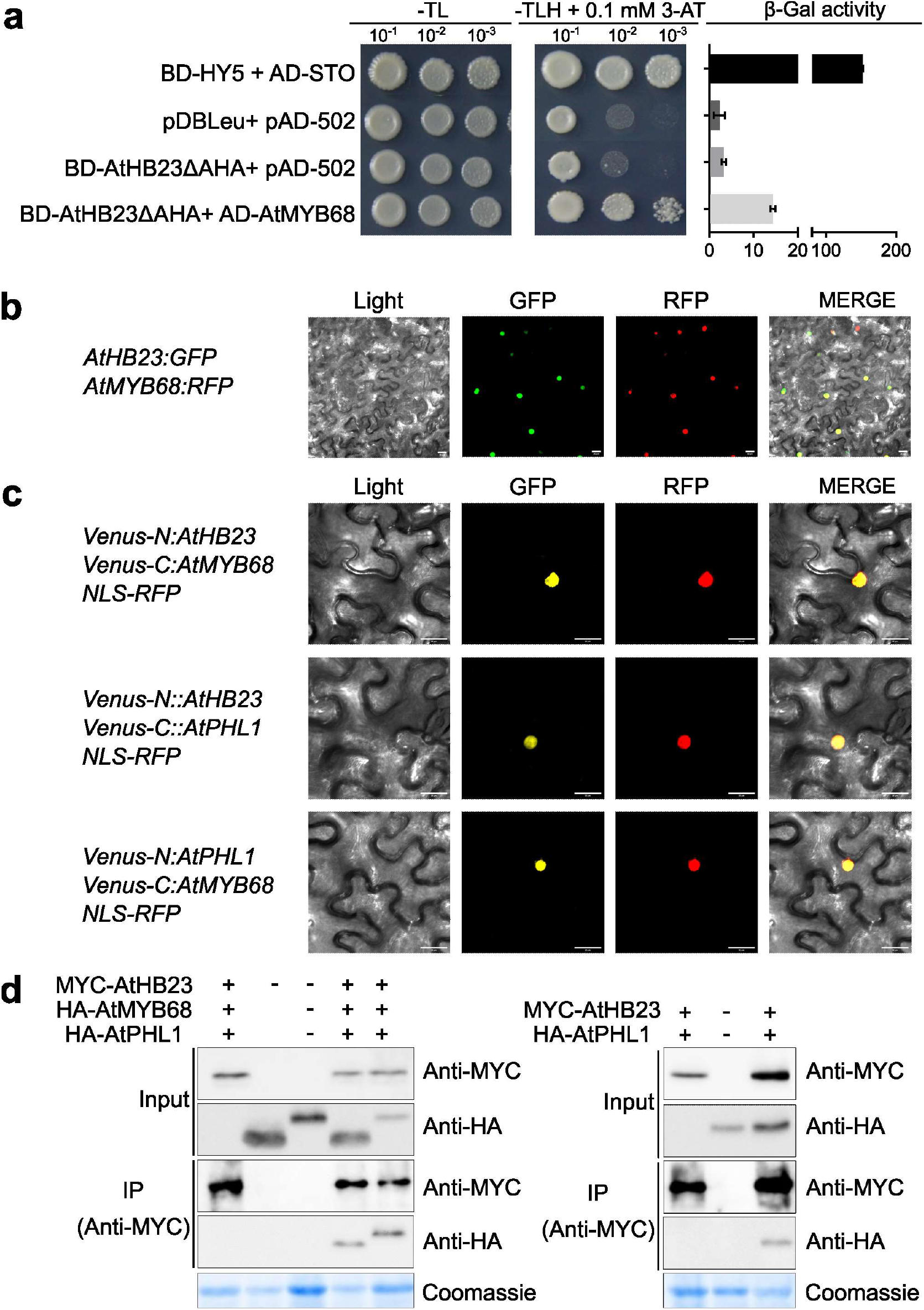
The transcription factors AtHB23, MYB68, and PHL1 interact with each other in yeast and *in planta*. **a**. Y2H assay performed with BD-AtHB23 ΔAHA + AD-MYB68 and the corresponding controls on the selection media (SD – Trp,Leu (-TL) and SD-Trp,Leu,His with 0.1 mM 3AT (-TLH+0.1mM 3-AT). On the right panel, β-galacosidase activity in miller units is shown. BD: Gal4 DNA binding domain; AD: Gal4 activation domain. BD-HY5 + AD-STO and pDBLeu + pAD-502 were used as positive and negative controls, respectively. AtHB23ΔAHA: the C-terminal AHA domain was removed (Capella *et al*., 2014). **b**. Colocalization assay using *Nicotiana benthamiana* leaves. Agrobacteria containing *35S:AtHB23:GFP* and *35S:AtMYB68:RFP* were infiltrated and pictures were taken 2 days after with a confocal microscope. GFP: GFP image, RFP: RFP image, Merge: merged fluorescent and light image. **c**. Mutual interactions between AtHB23, AtMYB68, and AtPHL1 *in planta*. Agrobacteria transformed with *N-YFP:AtHB23* and *C-YFP:AtMYB68*; *N-YFP:AtHB23* and *C-YFP:AtPHL1*; *N-YFP:AtPHL1*, and *C-YFP:AtMYB68* constructs were used. NLS-RFP was used to cotransform as nuclear localization signal. **d**. Co-immunoprecipitation assay *in vivo*. AtHB23 carrying an MYC tag (MYBC-HB23) was co-expressed with HA-tagged AtMYB68 (HA-AtMYB68) and AtPHL1 (HA-AtPHL1). The right panel shows the interaction between MYC-AtMYB68 and HA-AtPHL1. Altogether these results indicated that these three TFs interact with each other in all combinations in the plant nucleus.

Given that the established relationships were independent, AtHB23 with AtMYB68 or with AtPHL1, we wondered whether AtMYB68 and AtPHL1 interacted between them. Unfortunately, Y2H analysis between Gal4BD-AtHB68 and Gal4BD-AtPHL1 did not succeed because Gal4BD-AtMYB68 and Gal4BD-AtPHL1 showed strong transcription activities in yeast. Therefore, we performed a BiFC assay using VENUS-N:AtPHL1 and VENUS-C:AtMYB68 with positive results (Figure 1c and Supplementary Figure S1). Furthermore, these proteins colocalized in the nucleus when tobacco leaves were cotransformed (Figure 1c). A reciprocal combination of the three transcription factors constructed in the N-VENUS and C-VENUS vector sets resulted in crossed interactions between each other (Supplementary Figure S1). Next, we performed coimmunoprecipitation assays to confirm the obtained results by an independent method (Figure 1d). Total protein extracts from tobacco plants transiently transformed with MYC-AtHB23 plus HA-AtMYB68 and MYC-AtHB23 plus HA-AtPHL1 were pulled-down with anti-MYC antibodies (Figure 1d, left panel). Both pairs coimmunoprecipitated, supporting the above-described results (Figure 1d, right panel).

### *AtMYB68* and *AtPHL1* coexpress with *AtHB23* in specific tissues and developmental stages of root development

To test the functional sense of the interaction between these three TF, we first investigated the expression patterns of *AtMYB68* and *AtPHL1* in roots, the organ in which *AtHB23* was deeply investigated, associated with developmental and salinity responses (Perotti *et al*., 2019, 2020, and 2022). For this purpose, we used 8-day-old transgenic plants carrying the constructs *prAtMYB68:GUS* and *prAtPHL1:GUS*. *AtPHL1* was expressed in the root tip and the base of the lateral root primordium (LRP) in stages V to VII (Malamy and Benfey, 1997) and in the tip of emerged lateral roots (LR). GUS activity driven by the *AtMYB68* promoter was evident in the vascular tissue and throughout the developing LRP and LR (Figures 2 and Supplementary S2). *AtMYB68* and *AtPHL1* coincided with *AtHB23* at the base of LRP, hinting at a coordinated role of these interacting TF in LR development. Later, *AtMYB68* expression in LR development was restricted to the surrounding cells of the primordium, resembling that of other auxin-responsive genes involved in this developmental context (Figure 2; Marin *et al*., 2010).

**Figure 2.**
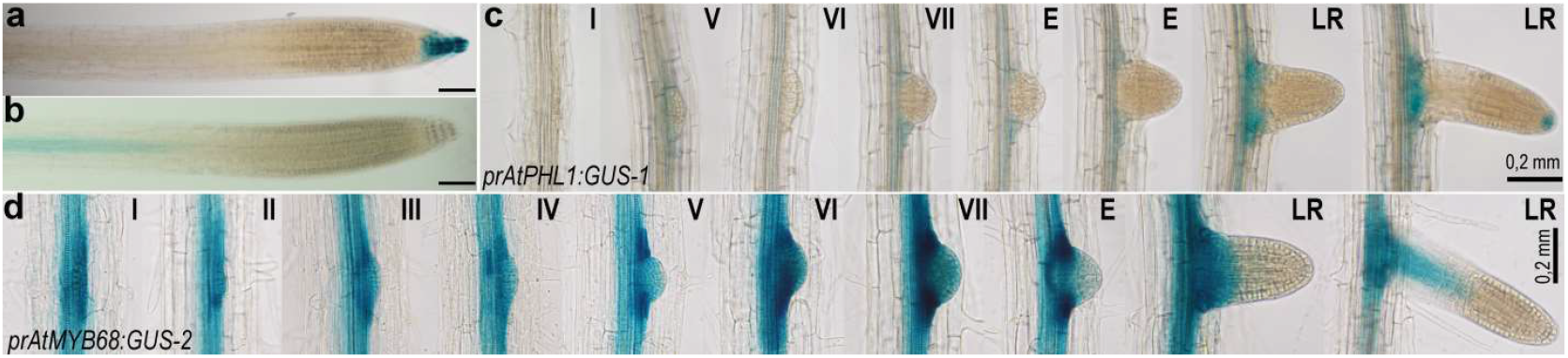
*AtMYB68* and *AtPHL1* genes are expressed in primary and lateral roots. Expression of *AtPHL1* (**a**) and *AtMYB68* (**b**) in the primary root, evaluated with *prAtPHL1:GUS* and *prAtMYB68:GUS* transgenic plants. Expression pattern of the same genes during lateral root development (**c** and **d**). I to VII represent different stages of lateral root primordium (LRP) and LR indicates emerged roots as described by Malamy and Benfey (1997). Black bar indicates 50 μm.

### AtMYB68 and AtPHL1 play a role in primary and lateral root development together with AtHB23

Considering tissue and subcellular colocalization of *AtMYB68*, *AtPHL1*, and *AtHB23*, plus their interaction in yeast and *in planta*, we investigated how these genes affect root architecture. For this purpose, we obtained *AtMYB68* silenced (*amiR68*; no mutants were available in the Col 0 background) and overexpressor (AT68) plants with altered transcript levels (Supplementary Figure S4). The number of initiated and emerged LRs in *amiR68* plants and *phl1* mutants was evaluated. *AtMYB68* silencing did not affect primary root length but significantly diminished LRP density (Figure 3a), the opposite phenotype to that shown by *amiR23* plants (Perotti *et al*., 2019). *Phl1* mutants and *amiR23* plants had longer primary roots, whereas *PHL1* overexpressors (ATPHL1) showed the opposite (Figure 3c). On the other hand, *phl1* mutants did not exhibit significant differences in the number of LRP or LR. *AtMYB68* overexpressors showed a similar scenario (Supplementary Figure S5).

**Figure 3.**
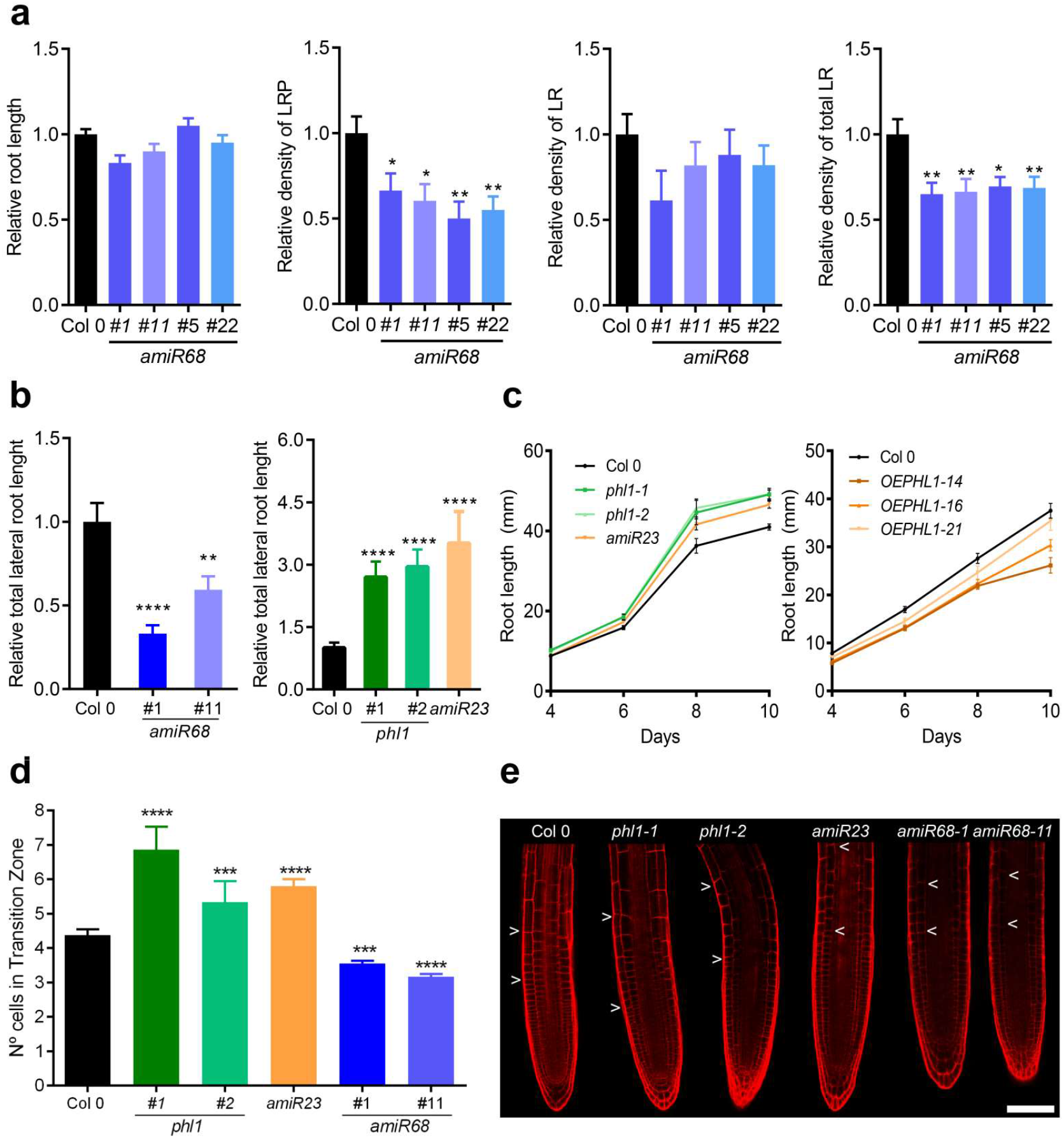
*AtMYB68* and *AtPHL1* modulate root architecture. **a**. Relative main root length in *amiR68* mutants compared with the Col 0 control. The relative density of LRP or LR was calculated as the number of LRP or LR/mm of the primary root and the relative density of total lateral roots (LRP + LR). The values were normalized with those measured in the control Col 0, taken as 1 (one). **b**. Total LR length of Col 0, *amiR68, phl1*, and *amiR23* mutants. **c**. Time course of root elongation of *phl1* and OEPHL1 relative to Col 0 plants. **d**. Number of cells in the transition zone of Col 0, *amiR68, phl1*, and *amiR23* mutants. **e**. Illustrative picture of confocal microscopy of root tips of Col 0, *amiR68, phl1* and *amiR23* mutants. Assays were repeated three times with N: 15/genotype Error bars represent SEM. Asterisks indicate significant differences doing a Student’s t-test (** P < 0.01, *** P < 0.001, **** P < 0.0001).

Remarkably, relative total lateral root length was diminished in *amiR68* plants and significantly augmented in *phl1* mutants, like in *amiR23* plants (Figure 3b). To test whether lateral root length differed between the three genotypes in cell number or cell size, LR root tips were analyzed by confocal microscopy. The analysis revealed fewer cells in *amiR68* plants in the transition zone, whereas *phl1* and *amiR23* mutant plants showed the opposite phenotype (Figures 3d and 3e).

Altogether, the results indicated a complex interplay between the three partners at the base of the LRP and the LR tip.

### Auxin induces *AtMYB68* expression impacting the hormone distribution in the root

Given that *AtHB23* expression in LR was regulated by auxin, and the encoded protein directly modulates the auxin carrier *LAX3*, we wondered whether its partners were also regulated by this hormone. *PrMYB68:GUS* plants were treated with 1 μM IAA and analyzed by histochemistry and RT-qPCR. Both assays indicated a strong induction of this gene by auxin in the root vascular system (Figures 4a and 4b). We also tested the effect of auxin on the expression in the root tip of *AtPHL1* and *AtHB23*. *AtPHL1* did not exhibit significant differences in the presence of IAA, whereas *AtHB23* showed a strong induction in the vascular system (Supplementary Figure S6). In view of the impact of IAA regulation on *AtMYB68*, we used *DR5:GUS* plants to cross them with *amiR68* mutants. Notably, *DR5* expression in the primary root tip disappeared in the crosses, indicating repression of the hormone transport to this tissue. The effect was similar in LR tips, whereas DR5 staining increased in the LRP (Figures 4c and 4d). To know the AtPHL1 influence, we generated new crosses between *DR5:GUS* and *phl1* mutants. In this case, the expression in the tips remained unaltered, whereas it disappeared from the vascular system (Figure 4e). Regarding auxin carriers, AUX1 appeared strongly induced by AtMYB68 because the crosses *prAUX1:GUS* × *amiR68* showed significantly less staining than *prAUX1:GUS* plants (Figures 4f and 4g).

**Figure 4.**
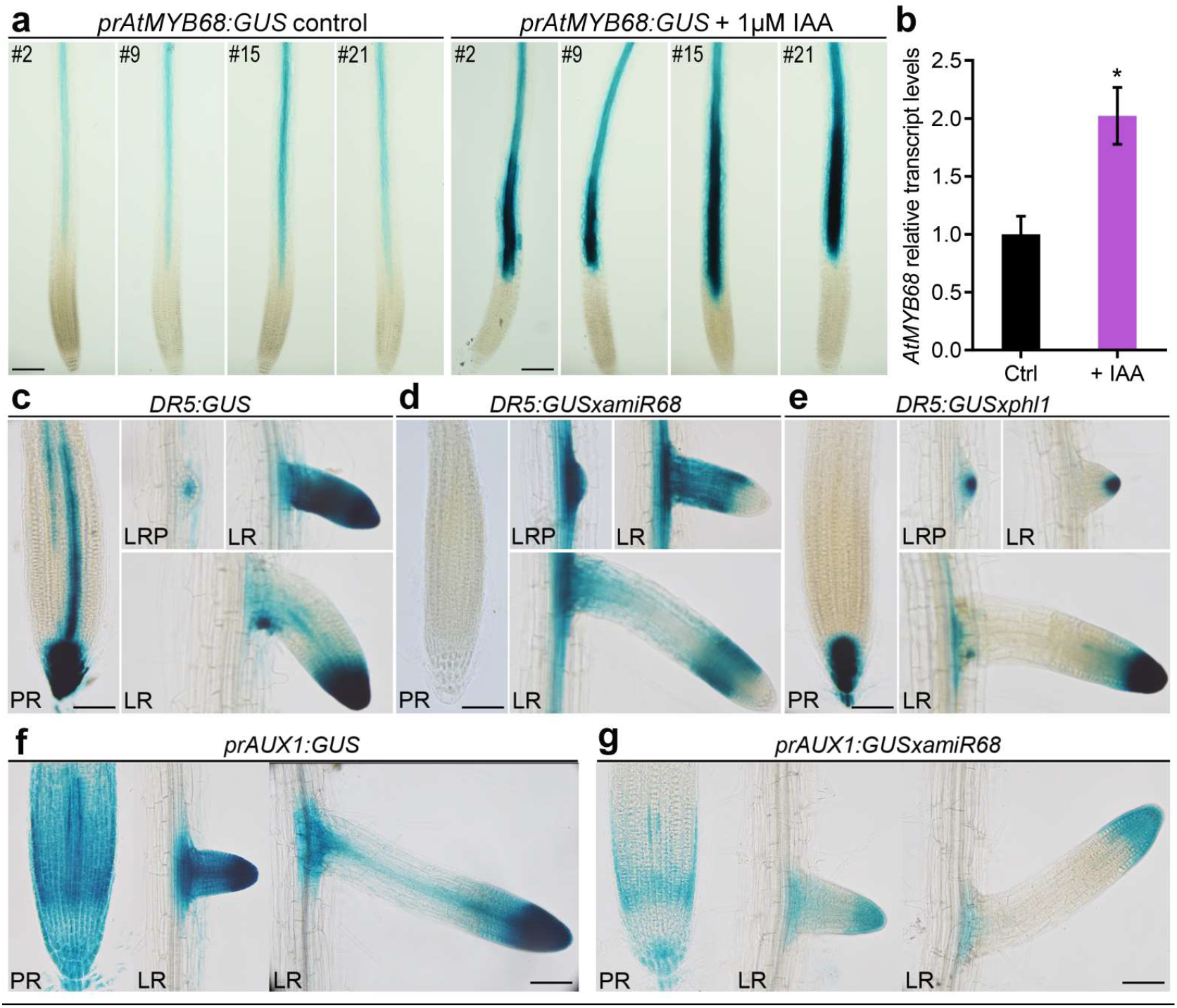
*AtMYB68* expression is induced by IAA and alters auxin distribution. **a**. GUS histochemistry of 8-day-old *prAtMYB68:GUS* roots (4 independent lines: *#2*, *#9*, *#15*, and *#21*) grown in control conditions (left panel) or treated with 1 μM IAA (right panel) for 12 h. The black bar indicates 50 μm. **b**. Transcript levels of *AtMYB68* in 7-day-old roots of seedlings grown in standard conditions or with 1 μM IAA added for 12 h. The value was normalized with that obtained in the control Col 0. The asterisk indicates a significant difference (*post hoc* Tukey test). GUS histochemistry of 8-day-old *DR5:GUS* (**c**), *DR5:GUS × amiR68* (**d**), *DR5:GUS × phl1* (**e**), *prAUX1:GUS* (**f**), and *prAUX1:GUS × amiR68* (**g**) roots. PR means primary root, LRP lateral root primordium, and LR emerged lateral roots. The black bar indicates 50 μm. Assays were repeated three times with N: 15/genotype.

### AtMYB68 or AtPHL1 play opposite roles in front of salinity conditions

Considering the impact of the interaction between AtMYB68, AtPHL1, and AtHB23 in root architecture and given the positive role of AtHB23 in salinity conditions, we wondered whether these MYB TFs were necessary for such a response. To answer this question, we analyzed the expression of these genes in salinity conditions. Transgenic plants carrying *prAtMYB68:GUS* and *prAtPHL1:GUS* were subjected to NaCl treatments and analyzed by histochemistry. *AtPHL1* was strongly induced in the root tip and the vascular system (Figure 5a), whereas *prAtMYB68:GUS* plants did not exhibit significant differences in GUS staining. However, *AtMYB68* transcript levels were significantly increased in Col 0 plants treated with NaCl (Figure 5b). To test whether the modulation of these genes’ expression by NaCl impacts mutant and overexpressor phenotypes, we counted survivors and dead plants after 9-15 days of treatment; *amiR68* and ATPHL1 plants showed similar sensitive phenotypes (Figure 5c). Moreover, ATPHL1 plants arrested the growth of the primary root, whereas *phl1* mutants showed the opposite behavior (Figure 5d). Considering total LR length, these mutants treated with NaCl resembled the Col 0 genotype, whereas *amiR68* and *amiR23* seedlings, which in normal conditions exhibited shorter LR, were less sensitive than the Col 0 under salinity considering this trait (Figure 5e).

**Figure 5.**
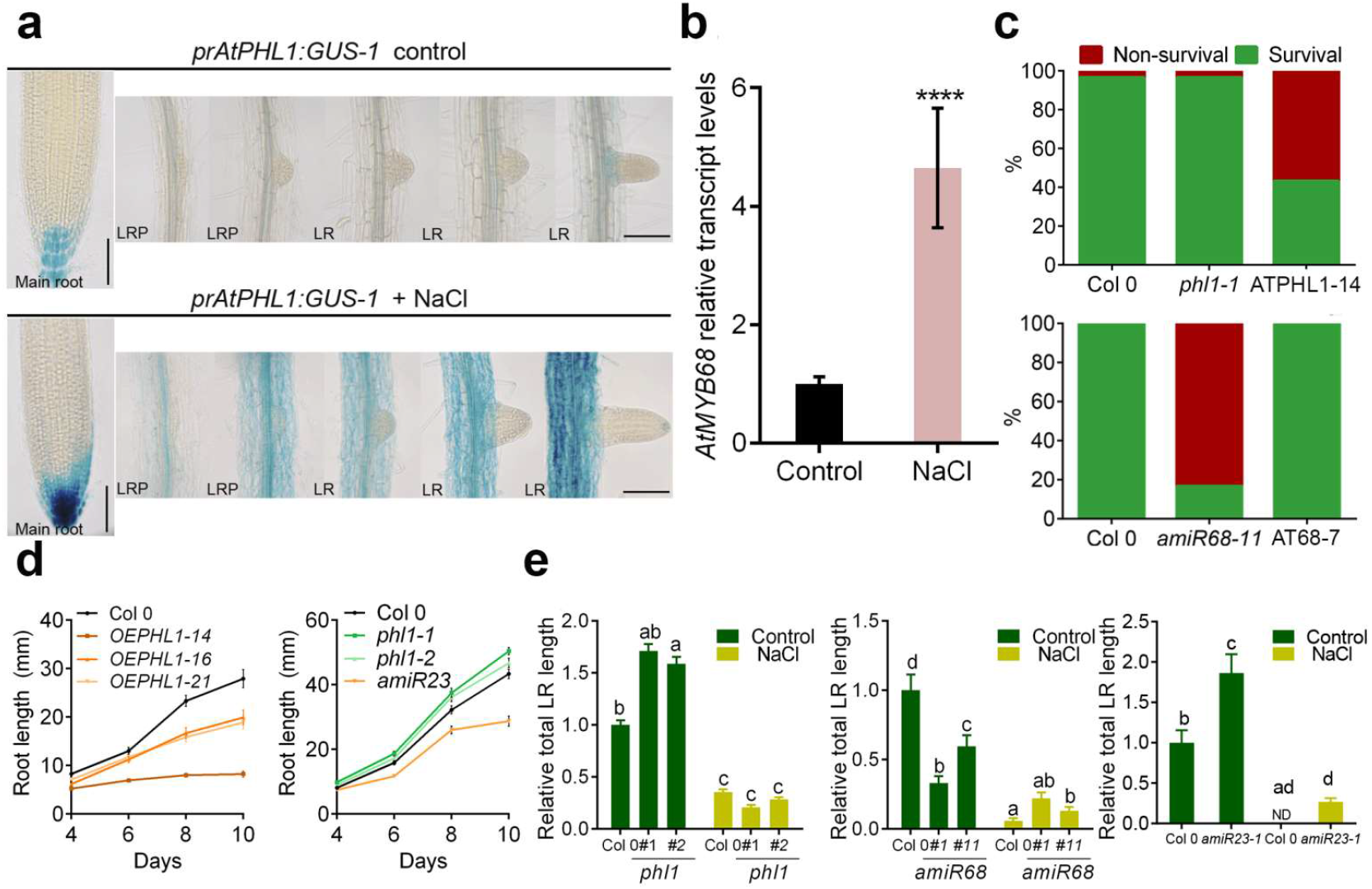
AtMYB68 has a positive role in front of salinity, whereas AtPHL1 is a negative regulator of such response. **a**. GUS histochemistry of 8-day-old *prAtPHL1:GUS-1* roots (upper panel) and after treatment with 100 mM NaCl (lower panel). **b**. Transcript levels of *AtMYB68* in 7-day-old roots of seedlings grown in standard conditions or with 100 mM NaCl for 12 h. The value was normalized with that obtained in the Col 0 control. The asterisk indicates a significant difference (*post hoc* Tukey test). **c**. Survival rate of plants (Col 0, *phl1*, ATPHL1, *amiR68*, and AT68) placed in plates with 100 mM NaCl, 3 days after sawing for 9-15 additional days. Red columns indicate the % of dead plants whereas green ones the % of survivors. **d**. Time course of the main root length evaluated in Col 0, OEPHL1 (three independent lines: *#14*, *#16*, and *#21*), *phl1* (two independent lines: −1, and −2) mutant, and *amiR23* seedlings grown in 75 mM NaCl. Quantitative measurements were performed from day 4 after sowing until day 10. **e**. Total LR length of Col 0, *amiR68, amiR23* and *phl1* mutants, grown in control conditions or treated with 75 mM NaCl. The assays were repeated at least three times with N: 15/genotype. The black bar represents 1 cm. Different letters indicate significant differences (Tukey test, P < 0.01). Error bars represent SEM.

### The adaptation ability of *AtHB23*, *AtPHL1*, and *AtMYB68* mutants, overexpressors, and crossed plants to salinity correlates with the starch granules state in the root tip

Root gravitropism depends on the auxin gradient between the upper and lower sides (Zhang *et al*., 2019). In the columella cells, starch aggregates are formed, named statoliths or amyloplasts (Leitz *et al*., 2009). A salinity medium severely affects the auxin gradient and consequently, amyloplast formation. *AtHB23*, *AtPHL1*, and *AtMYB68* mutant and overexpressor plants differentially responded in front of salinity stress (Figure 5). To understand this scenario, we analyzed starch content staining the root tips of these plants with a Lugol solution (Figure 6). Seedlings were grown in normal conditions for five days (Figure 6a) and then placed in 150 mM NaCl for seven-eight hours (Figure 6b). It was previously shown that *amiR23* plants significantly reduced their starch content after this treatment (Perotti *et al*., 2022 and Figure 6b). As expected, *amiR68* and ATPHL1 genotypes exhibited the same aspect (Figure 6b). The phenotype of *amiR68* seedlings was rescued in plants crossed with the AT68 genotype, indicating that the silencing of this gene was responsible for the NaCl-enhanced sensitivity (Figure 6b). Afterward, one-half of the seedlings were transferred back to MS medium while the other half remained in NaCl. Like AT23 plants, *phl1*, and AT68 roots slowly adapted to the salinity medium and recovered their starch granules as they did in the NaCl-free medium (Figures 6c and 6d), whereas *amiR68* and ATPHL1 genotypes were unable to restore a healthy phenotype (Figures 6c, 6d, and Supplementary S7).

**Figure 6.**
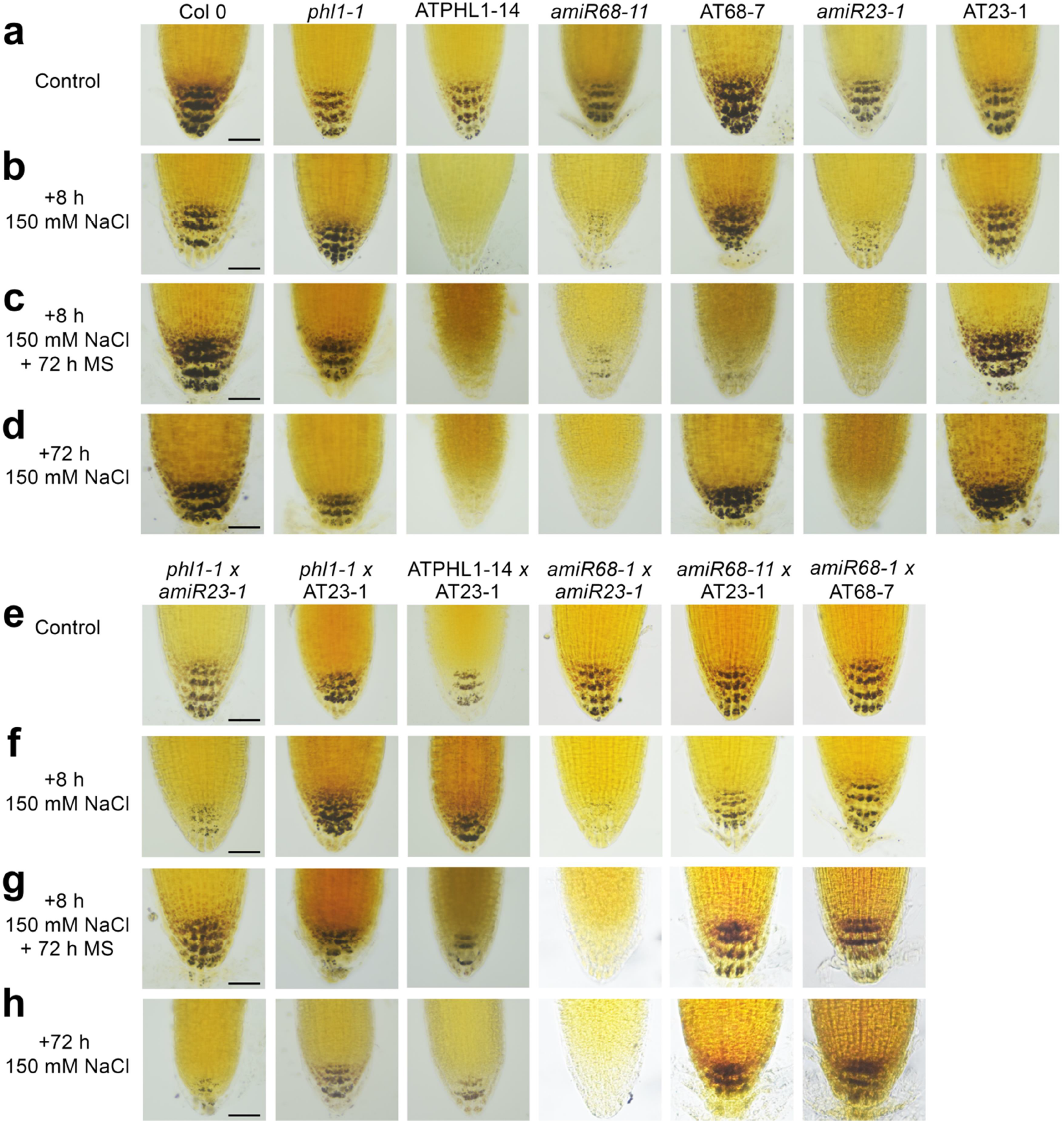
The adaptation ability to salinity depending on *AtHB23*, *AtPHL1*, and *AtMYB68* levels correlates with the starch granules stage in the root tip. **a**. Illustrative pictures of root tips (5-day-old) stained with Lugol solution of Col 0, *phl1*, OEPHL1, *amiR68*, AT68, *amiR23*, and AT23 seedlings grown in normal conditions. **b**. After 8 h treatment with 150 mM NaCl. **c**. The roots were transferred to normal conditions or maintained in 150 mM NaCl for additional 72 h (**d**). **e-h**. The same analysis was carried out with the crossed plants *phl1* × *amiR23*, *phl1* × AT23, OEPHL1 × AT23, *amiR68* × *amiR23*, and *amiR68* × AT23. **e**. Control conditions. **f**. after 8 h in 150 mM NaCl. **g**. after 8 h 150 mM NaCl + 72 h in MS. **h**. after 8 h + 72 h in 150 mM NaCl. The black bar represents 50 μm.

Given these results, we stated the hypothesis based on the availability of AtHB23 to exert a positive action dealing with salinity, avoided by AtPHL1 or enhanced by AtMYB68. To test this, we obtained crossed plants and assayed their behavior by performing the same assay described above. *AmiR23* × *amir68* plants lost their amyloplasts and could not recover them even after 72 h in normal conditions (Figures 6e-h). Notably, *amiR68* × AT23-crossed plants did not lose their starch granules after the NaCl treatment, suggesting that the overexpression of *AtHB23* compensated somehow for the low availability of AtMYB68 to cooperatively interact. On the other hand, *phl1-1 × amiR23* seedlings behaved like the *amiR23* genotype, supporting the essential role of AtHB23 in the positive response in front of salinity. In ATPHL1 × AT23 crosses, the picture was intermediate between that of the parent genotypes (Figures 6e-h).

Regarding these results, we evaluated transcript levels of genes encoding key enzymes participating in starch synthesis and degradation in *phl1* and *amiR68* plants. *ADG1* and *PGM*, involved in starch synthesis, did not significantly change, except in *amiR68* roots, where they were slightly reduced in salinity and after recovery, respectively (Supplementary Figure S8). *BAM1*, participating in degradation, was induced in *amiR68* plants in salinity, whereas *phl1* mutants behaved similarly to the WT, whereas GWD did not show differences. These results indicated that starch turnover was altered by the levels of *AtMYB68* and *AtPHL1* (Supplementary Figure S8). However, they alone cannot explain the absolute lack of starch observed in *amiR68* plants and the amyloplast integrity in *phl1* mutants, indicating that other mechanisms must be modulating this process.

## DISCUSSION

Root plasticity is crucial for plant adaptation to soil and involves the growth or arrest of the primary, lateral, and high-order roots. These events are fine-modulated by many biomolecules, such as transcription factors (TF) and phytohormones. Many detailed studies reported the functional characterization of TF in lateral root initiation, emergence, and elongation, as well as in primary root growth (Banda *et al*., 2019). These studies were performed in normal growth conditions and under different stress factors (Ambastha *et al*., 2020; Verma *et al*., 2022). However, the involvement of an individual TF in the global architecture determination, including primary and lateral roots, is less abundant. Here, we reported how three TF, one HD-Zip, and two MYB family members interact to activate or repress primary and lateral root development, depending on the environmental condition.

AtHB23, the most studied among the three proteins, was shown to be non-redundant with its putative paralogue AtHB13 in roots (Perotti *et al*., 2019). AtPHL1 was studied only in front of Pi starvation in a secondary role compared with its paralogue AtPHR1 (Bustos *et al*., 2010). *AtMYB68* was expressed in the root pericycle of the Ler ecotype, responding to environmental cues (Feng *et al*., 2004), and detected in the Arabidopsis root protein expression landscape (Petricka *et al*., 2012). It was also studied in the reproductive stage, determining seed yield and tolerance to drought and high temperatures (Deng *et al*., 2020).

Several members of the HD-Zip I family are expressed in different root tissues (Perotti *et al*., 2021), and a few were functionally characterized as involved in root development and response to stress (Miao *et al*., 2018; Mora *et al*., 2022). MYB-CC proteins were studied in several species, particularly associated with Pi starvation (Bai *et al*., 2019; Bhutia *et al*., 2020). The large MYB family has many well-characterized members acting in roots. For example, AtMYB77 was shown to regulate a subset of auxin-responsive genes during LR development and interacted *in vitro* with ARF proteins. The knockout mutant *atmyb77* exhibits a lower density of LRs than the WT (Shin *et al*., 2007). Also, AtMYB93 is an auxin-inducible gene acting as a negative regulator of LR development in Arabidopsis, making part of an auxin-triggered negative feedback loop, ensuring that lateral roots only develop when required (Gibbs *et al*., 2014). Finally, AtMYB36 was reported as a regulator of the transition from proliferation to differentiation in the endodermis. The characterization of *atmyb36* mutants suggested that this TF acts as a positive regulator of differentiation and a negative regulator of proliferation in root meristems (Liberman *et al*., 2015).

Although there are significant differences in root development depending on the ecotype (Perotti *et al*., 2020), the expression pattern reported in Ler plants (Feng *et al*., 2012) was similar to the informed in this manuscript.

There is abundant literature about TF transcriptional, post-transcriptional, and post-translational regulation, influencing their stability or activity (Deribe *et al*., 2010; Nelson and Millar, 2015; Zhang *et al*., 2021; Zhu, 2016). However, it is hard to find literature describing TF functioning in both primary and LR development and also about the association between members of different families acting cooperatively or in opposite ways by protein-protein associations. The TF partitioning between the nucleus and the cytoplasm is an essential mechanism regulating developmental events and adaptation (Allen and Strader, 2021). Such is the case of the interaction between the HD-Zip I TF HaHB11 from *Helianthus annuus* and AtHB7 from Arabidopsis, modulated by the Kinesin13B (Miguel *et al*., 2020). Even scarce, it was recently informed that the module OsFTIP6-OsHB22-OsMYBR57 modulates drought tolerance in rice (Yang *et al*., 2022). Notably, OsHB22 is an HD-Zip I TF previously reported as a negative regulator in ABA-mediated drought and salt tolerances in rice (Zhang *et al*., 2012). Rice mutants in this gene behaved better under drought stress showing no yield penalty (Zhang *et al*., 2012). OsMYBR57 is an MYB-related protein; its mutant displayed a drought-sensitive phenotype. Yang *et al*. (2022) revealed that it interacts with the HD-Zip OsHB22 and both TF together modulate the expression of several bZIP TF participating in the drought response. Another example is the interaction in the nucleus between XIW1 (XPO1-interacting WD40 protein 1) and ABI5 (ABA INSENSITIVE 5), modulating ABA response (Xu *et al*., 2019).

Coexpression of TF in the same tissue, developmental stage, and environmental condition is an absolute requirement to suggest interaction. We showed that *AtHB23*, *AtMYB68*, and *AtPHL1* are expressed in the primary root tip and specific stages of LR development (Figure 2). Remarkably, *amiR68* mutants exhibited fewer LRP, the opposite phenotype of *amiR23* plants (this paper and Perotti *et al*., 2019), suggesting that AtHB23 needs AtMYB68 to exert its function. AtPHL1 does not participate in LR initiation (Supplementary Figure S5) but in LR elongation, having a cooperative role with AtHB23, opposing that of AtMYB68, at least in normal conditions (Figure 3). Regarding primary root growth, AtMYB68 seems absent, whereas AtPHL1 and AtHB23 exhibit opposite roles (Figure 3).

AtMYB68 was induced by auxin in the vascular system, and although we could not detect its expression in the root tip of *prAtMYB68:GUS* seedlings, it seriously affected auxin distribution in this tissue, as shown by the crosses *DR5:GUS* × *amiR68* (Figure 4). AtPHL1 did not affect auxin in the root tip of primary and LR but in the vascular system. Among auxin carriers, AUX1 was involved in LR initiation and LAX3 in LR emergence (Marchant *et al*., 1999, 2002). AtHB23 regulated *LAX3*, whereas AtMYB68 modulated *AUX1* expression in the primary and LR.

The three TF are induced in salinity conditions exerting cooperative (AtHB23 and AtMYB68), and opposite (AtPHL1) functions (Figure 5). Like *amiR23*, *amiR68*, and ATPHL1 plants showed a reduced survival capacity in 150 mM NaCl, accompanied by a fewer ability to elongate primary roots exploring a less saline medium. Regarding LR elongation, *amiR68* plants were less penalized in NaCl than in control conditions, whereas *phl1* mutants lost their more elongated phenotype.

Under abiotic stress conditions, the survival ability correlated with the conservation or degradation of amyloplasts in the columella cells (Takahashi *et al*., 2003). Notably, LR overlap salinity conditions that are lethal for the primary root (Ambastha *et al*., 2020). *AtHB23* silencing provoked the loss of starch granules (Perotti *et al*., 2022). After analyzing amyloplasts in single mutants and crosses, we propose that since AtHB23 is necessary to deal with salinity and silenced plants cannot survive in such conditions, the action of this gene is fine-tuned. The phenotype under salinity of the crossed plants supported this interpretation (Figure 6). AtMYB68 interaction is required for this function, and AtPHL1 kidnaps both TF by protein-protein interactions to modulate such response (Figure 7).

**Figure 7.**
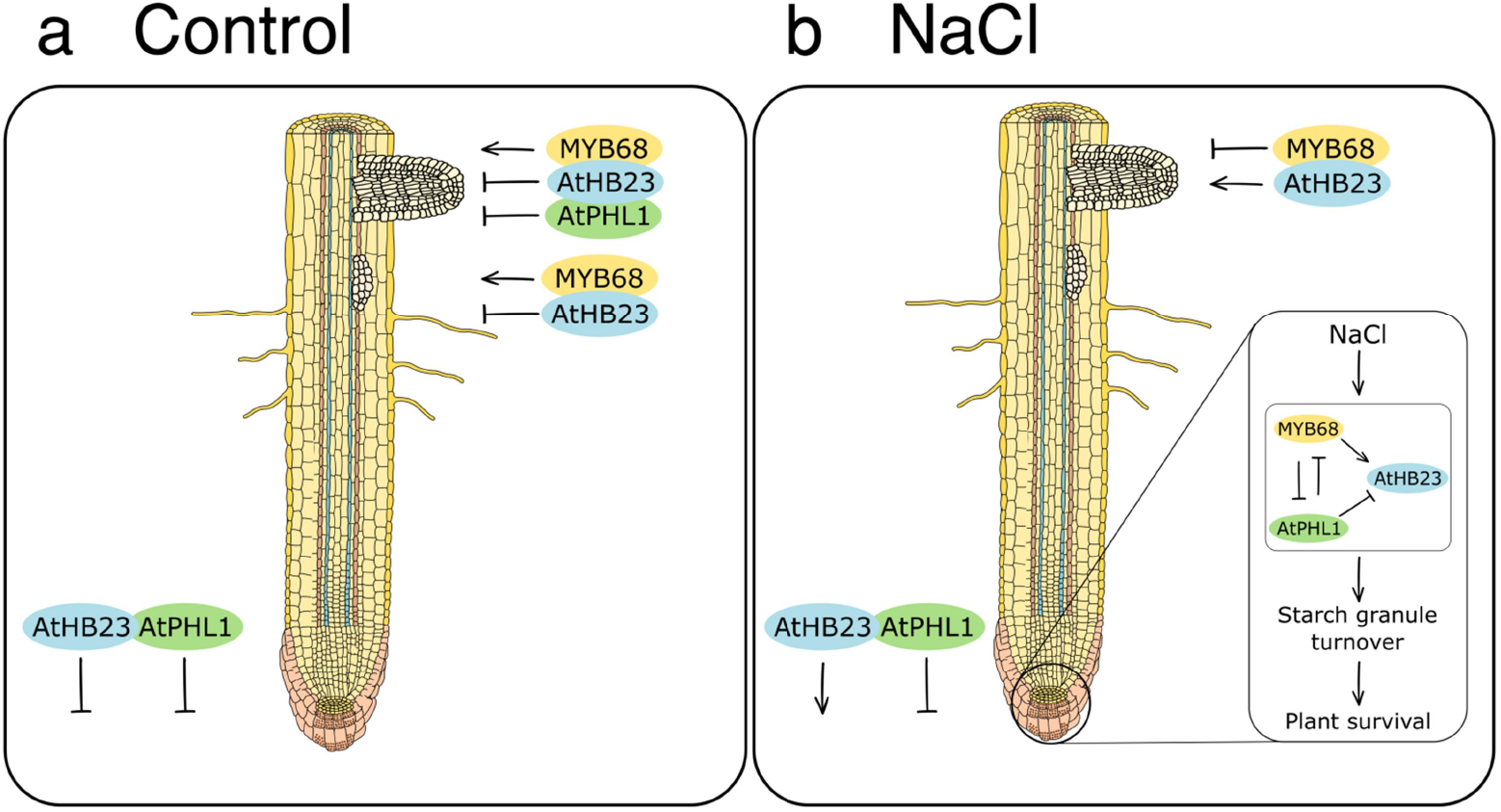
Proposed model for the interactive action of AtHB23, AtPHL1, and AtMYB68 in normal and salinity conditions. Proposed model for AtHB23, AtPHL1, and AtMYB68 roles in the primary and lateral root development in control (**a**) and salinity (**b**) conditions. Arrows between actors indicate direct regulation.

Altogether, our results indicated a fine regulation of primary and LR development in normal growth conditions and dealing with salinity by the interplay between AtHB23, AtMYB68, and AtPHL1 (Figure 7).

## Supporting information

Moreover, their interaction was confirmed by BiFC assays in Nicotiana benthamiana leaves (Figure 1c and Supplementary Figure S1).

## ABBREVIATIONS

HD-Zip: homeodomain-leucine zipper
TF: transcription factor
WT: wild type
GUS: β-Glucuronidase
Col 0: Columbia ecotype

## SUPPLEMENTARY MATERIAL

**Figure S1. The transcription factors AtHB23, AtMYB68, and AtPHL1 interact with each other in all the combinations**

BiFC analysis of protein-protein interaction between AtHB23, AtMYB68, and AtPHL1 using agroinfiltrated *Nicotiana benthamiana* leaves. The six combinations of N-terminal and C-terminal YFP; Venus-N for N-YFP and Venus-C for C-YFP, respectively, are shown. Positive control was carried out with HY5 and STO, and negative one using the empty vector.

**Figure S2. *AtMYB68* is expressed during root development**

Expression of *AtMYB68* independent lines in the primary root, evaluated with *prAtMYB68:GUS* transgenic plants. Expression pattern during lateral root development. I to VII represent different stages of lateral root primordium (LRP) and LR indicates emerged roots as described by Malamy and Benfey (1997). Black bar indicates 50 μm.

**Figure S3. *AtPHL1* is expressed during root development**

Expression of *AtPHL1* using the independent line *prAtPHL1-2:GUS* in the primary root and lateral root, evaluated using *prAtPHL1-2:GUS* transgenic plants. Black bars indicate 50 μm.

**Figure S4. Relative transcript levels of *AtMYB68* in overexpressor and silenced plants**

Transcript levels of *AtMYB68* in 10-day-old seedlings grown in normal conditions of AT68 and *amiR68* genotypes. The values were normalized with that obtained in Col 0. Bars represent SEM. Data were analyzed using a two-way ANOVA considering genotype and treatment. Different letters indicate significant differences (Tukey test, P < 0.01).

**Figure S5. AT68 and *phl1* mutant plants did not exhibit differential root phenotypes**

**a.** Relative main root length in Col 0 and two independent transgenic ATMYB68 lines (AT68: *#*5 and *#*7). The relative density of LRP or LR was calculated as the number of LRP or LR/mm of primary root and the relative density of total lateral roots (LRP + LR). The values were normalized with those measured in the Col 0 control, taken as 1 (one). **b.** Relative main root length in Col 0 and two *phl1* mutants (*#*1 and *#*2). The relative density of LRP or LR was calculated as the number of LRP or LR/mm of the main primary root and the relative density of total lateral roots (LRP + LR).

Assays were repeated three times with N: 15/genotype Error bars represent SEM. Different letters indicate significant differences (Tukey test, P < 0.01).

**Figure S6. *AtHB23* expression is induced by IAA in primary root, whereas that of AtPHL1 did not**

GUS histochemistry of 8-day-old *prAtAtHB23_L_:GUS* (**a**) and *prAtPHL1:GUS-1* (**b**) roots (3 independent lines, *#1*, *#2*, and *#3*) grown in control conditions or treated with 1 μM IAA during 12 h. The black bar indicates 50 μm.

**Figure S7. The adaptation ability to salinity depending on *AtHB23*, *AtPHL1*, and *AtMYB68* levels correlates with the starch granules stage in the root tip**

**a**. Illustrative pictures of root tips (5-day-old) stained with Lugol solution of Col 0, *phl1-2*, OEPHL1-16, *amiR68-5*, and AT68-5 seedlings grown in normal conditions. **b**. The same genotypes after 8 h treatment with 150 mM NaCl. **c**. The roots were transferred to normal conditions or maintained in 150 mM NaCl for additional 72 h (**d**). The black bar represents 50 μm.

**Figure S8. Starch synthesis and degradation are affected by AtPHL1 and AtMYB68**

Transcript levels of key genes in WT (Col 0), *phl1-1*, and *amiR68–5* plants grown in normal conditions for 5 days treated 7 h with 150 mM NaCl, and placed to recover in MS medium for additional 72 h. Genes assessed participating from degradation were *GWD* and *BAM1* (**a**), and from synthesis *PGM* and *ADG1* (**b**). All the values were normalized with the one obtained in the WT (Col 0). Bars represent SEM. Data were analyzed using a two-way ANOVA considering genotype and treatment. Different letters indicate significant differences (Tukey test, P < 0.01).

## ACKNOWLEDGEMENTS

This work was supported by Agencia Nacional de Promoción Científica y Tecnológica (PICT 2017 0305, PICT 2019 01916, and PICT 2020 0805 to RLC), and CONICET, grants from the Basic Science Research Program through the National Research Foundation of Korea (NRF), funded by the Ministry of Education (2020R1A6A1A03044344 and 2020R1F1A1074027) to JCH. FPS and MFP are CONICET Ph. D. Fellows. RLC is a CONICET Career member.

We thank to Dr. Ranjan Swarup for kindly providing *AUX/LAX* promoters fused to *GUS* seeds used in this study to our colleague Dr. Javier Moreno.

## AUTHOR CONTRIBUTIONS

FPS and MFP carried out most experiments and the corresponding figures. CIJ and JCH performed the TF library, screened it with AtHB23, and corroborated the results by Y2H and BiFC analyses. RLC conceived, designed, and wrote the manuscript. All the authors carefully revised and approved the manuscript.

