## Supplementary material for "A complex tissue-specific interplay between the Arabidopsis transcription factors AtMYB68, AtHB23, and AtPHL1 modulates primary and lateral root development and adaptation to salinity": Moreover, their interaction was confirmed by BiFC assays in Nicotiana benthamiana leaves (Figure 1c and Supplementary Figure S1).

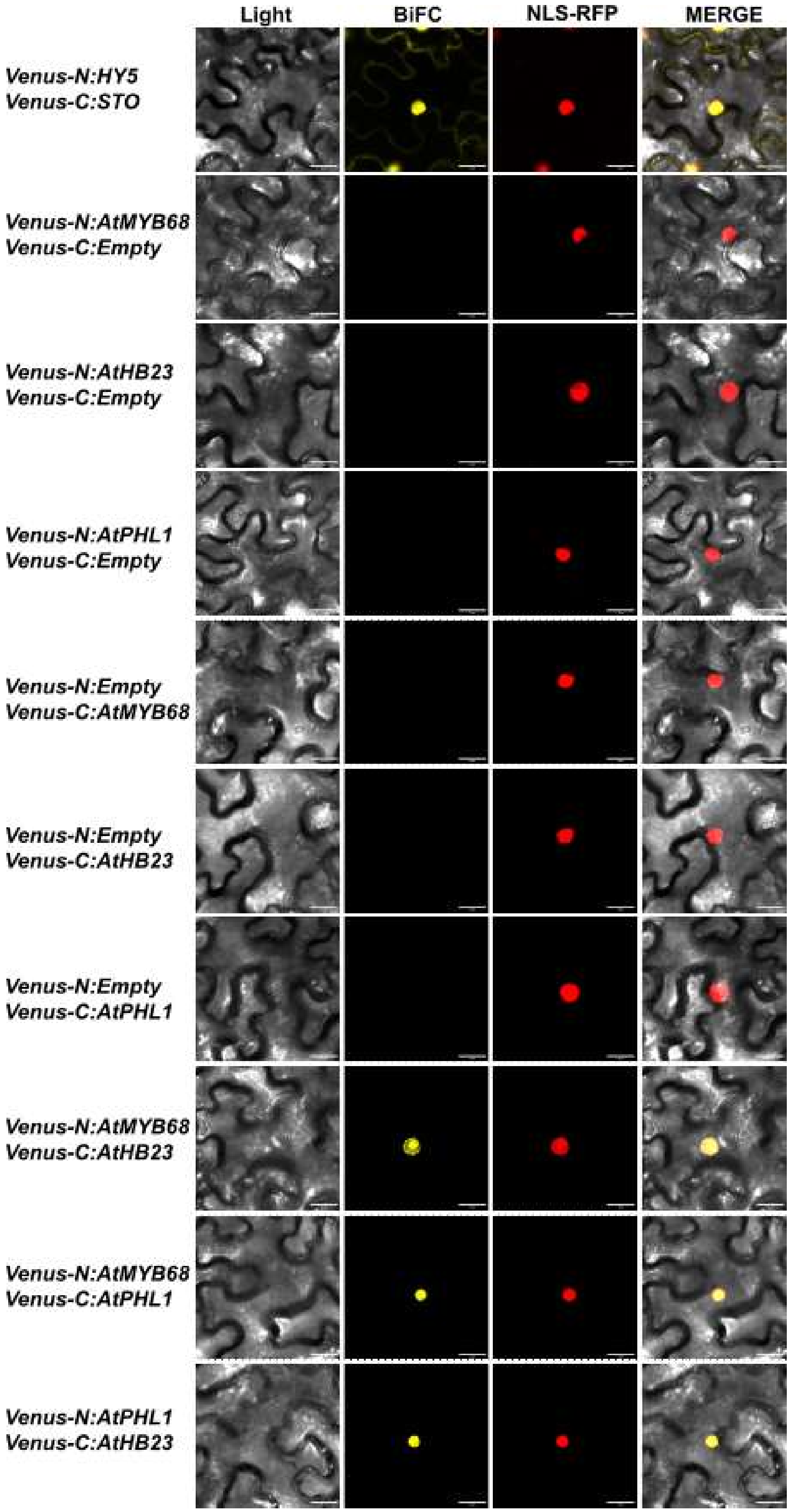
Light
BiFC
NLS-RFP
MERGE
Venus-N:HY5
Venus-C:STO
Venus-N:AtMYB68
Venus-C:Empty
Venus-N:AtHB23
Venus-C:Empty
Venus-N:AtPHL1
Venus-C:Empty
Venus-N:Empty
Venus-C:AtMYB68
Venus-N:Empty
Venus-C:AtHB23
Venus-N:Empty
Venus-C:AtPHL1
Venus-N:AtMYB68
Venus-C:AtHB23
Venus-N:AtMYB68
Venus-C:AtPHL1
Venus-N:AtPHL1
Venus-C:AtHB23

**Figure S1**. **The transcription factors AtHB23, AtMYB68, and AtPHL1 interact with each other in all the combinations**

BiFC analysis of protein-protein interaction between AtHB23, AtMYB68, and AtPHL1 using agroinfiltrated *Nicotiana benthamiana* leaves. The six combinations of N-terminal and C-terminal YFP; Venus-N for N-YFP and Venus-C for C-YFP, respectively, are shown. Positive control was carried out with HY5 and STO, and negative one using the empty vector.

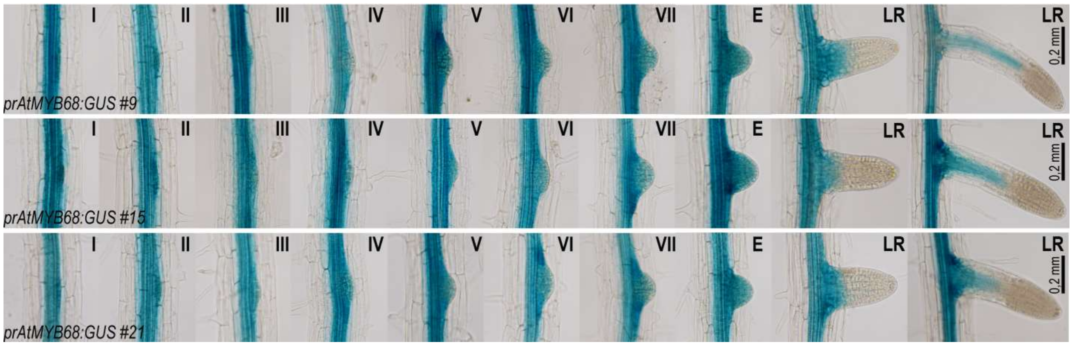


**Figure S2. *AtMYB68* is expressed during root development**

Expression of *AtMYB68* independent lines in the primary root, evaluated with *prAtMYB68:GUS* transgenic plants. Expression pattern during lateral root development. I to VII represent different stages of lateral root primordium (LRP) and LR indicates emerged roots as described by Malamy and Benfey (1997). Black bar indicates 50 µm.

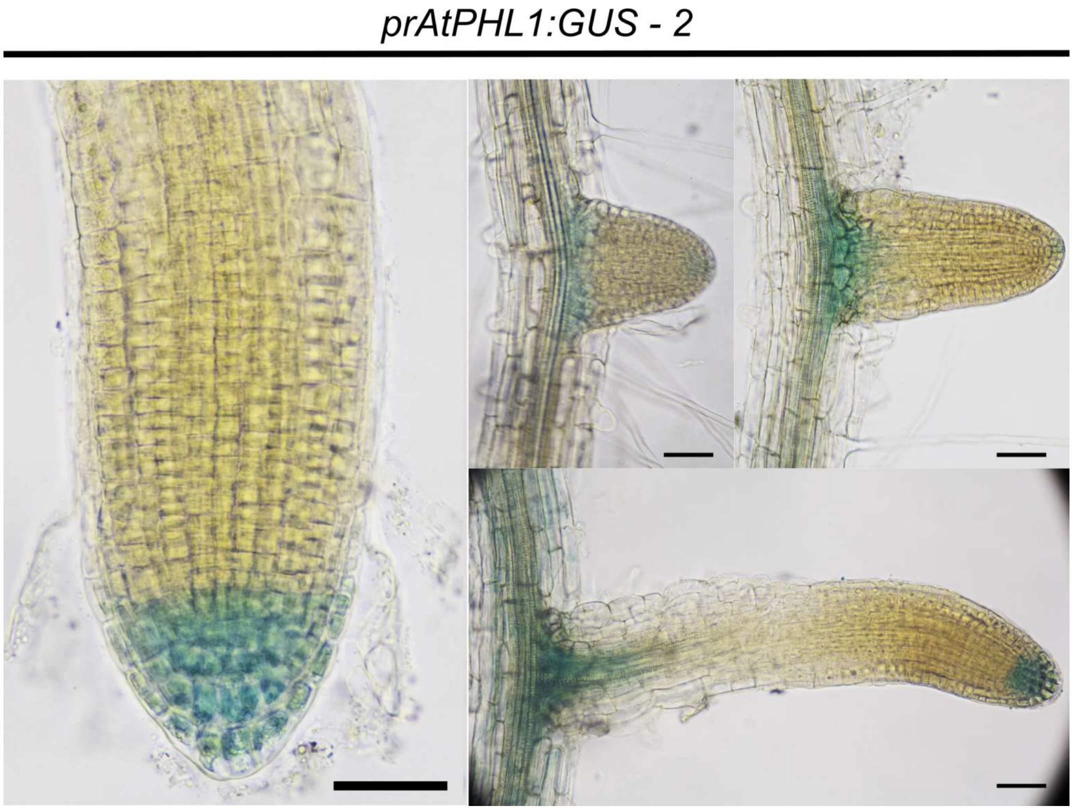


**Figure S3. *AtPHL1* is expressed during root development**

Expression of *AtPHL1* using the independent line *prAtPHL1-2:GUS* in the primary root and lateral root, evaluated using *prAtPHL1-2:GUS* transgenic plants. Black bars indicate 50 µm.

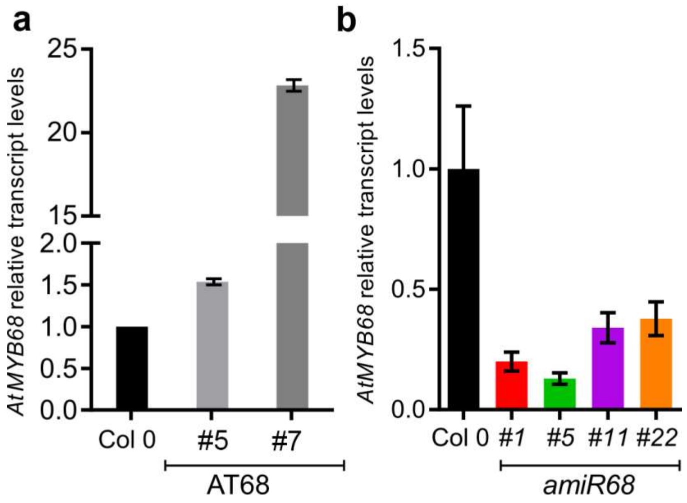


**Figure S4**. **Relative transcript levels of *AtMYB68* in overexpressor and silenced plants**

Transcript levels of *AtMYB68* in 10-day-old seedlings grown in normal conditions of AT68 and *amiR68* genotypes. The values were normalized with that obtained in Col 0. Bars represent SEM. Data were analyzed using a two-way ANOVA considering genotype and treatment. Different letters indicate significant differences (Tukey test, $P < 0.01$).

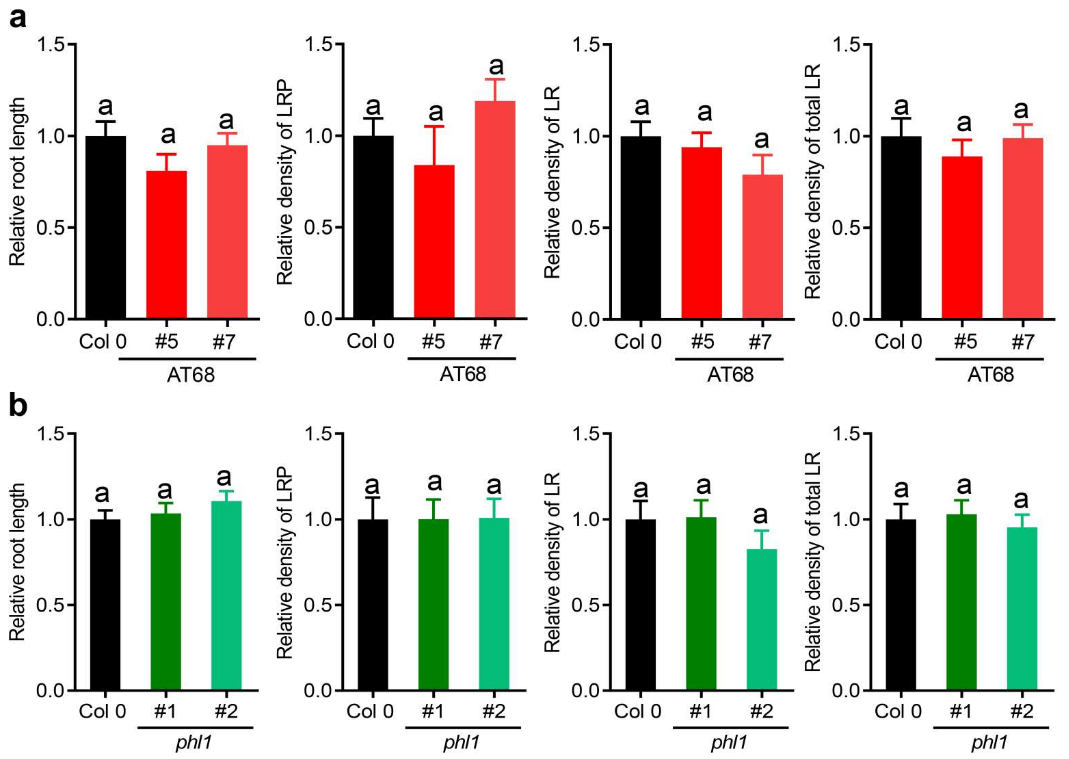


**Figure S5. AT68 and *phl1* mutant plants did not exhibit differential root phenotypes**

**a.** Relative main root length in Col 0 and two independent transgenic ATMYB68 lines (AT68: #5 and #7). The relative density of LRP or LR was calculated as the number of LRP or LR/mm of primary root and the relative density of total lateral roots (LRP + LR). The values were normalized with those measured in the Col 0 control, taken as 1 (one).

**b.** Relative main root length in Col 0 and two *phl1* mutants (#1 and #2*)*. The relative density of LRP or LR was calculated as the number of LRP or LR/mm of the main primary root and the relative density of total lateral roots (LRP + LR).

Assays were repeated three times with N: 15/genotype Error bars represent SEM. Different letters indicate significant differences (Tukey test, $P < 0.01$).

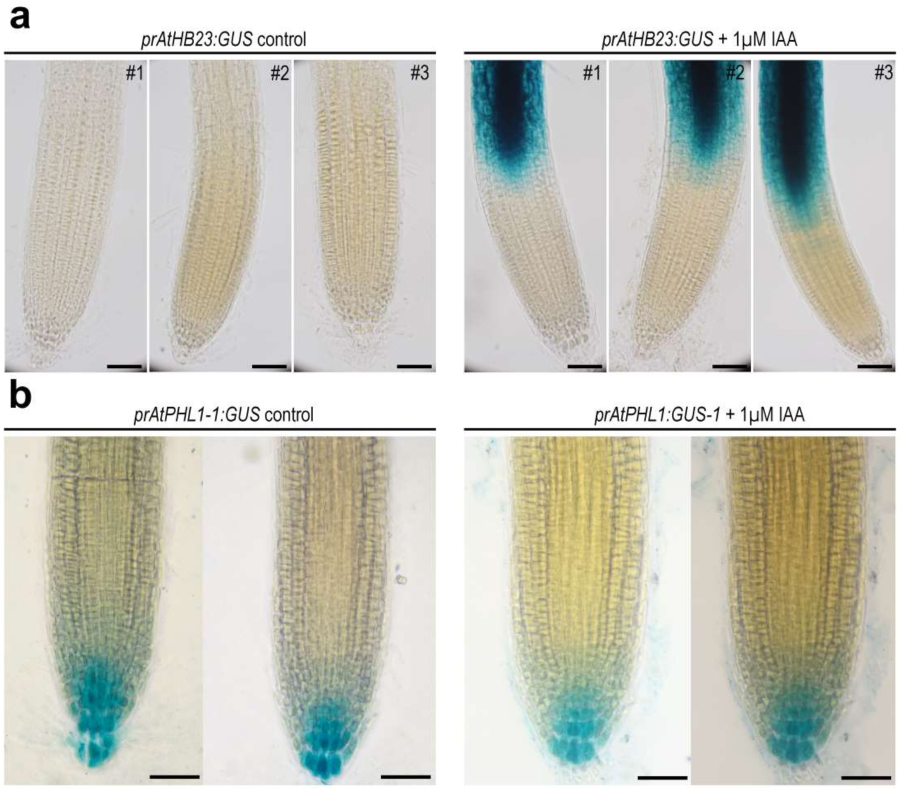


**Figure S6. *AtHB23* expression is induced by IAA in primary root, whereas that of AtPHL1 did not**

GUS histochemistry of 8-day-old *prAtAtHB23$_L$:GUS* (**a**) and *prAtPHL1:GUS-1* (**b**) roots (3 independent lines, *#1*, *#2*, and *#3*) grown in control conditions or treated with 1 µM IAA during 12 h. The black bar indicates 50 µm.

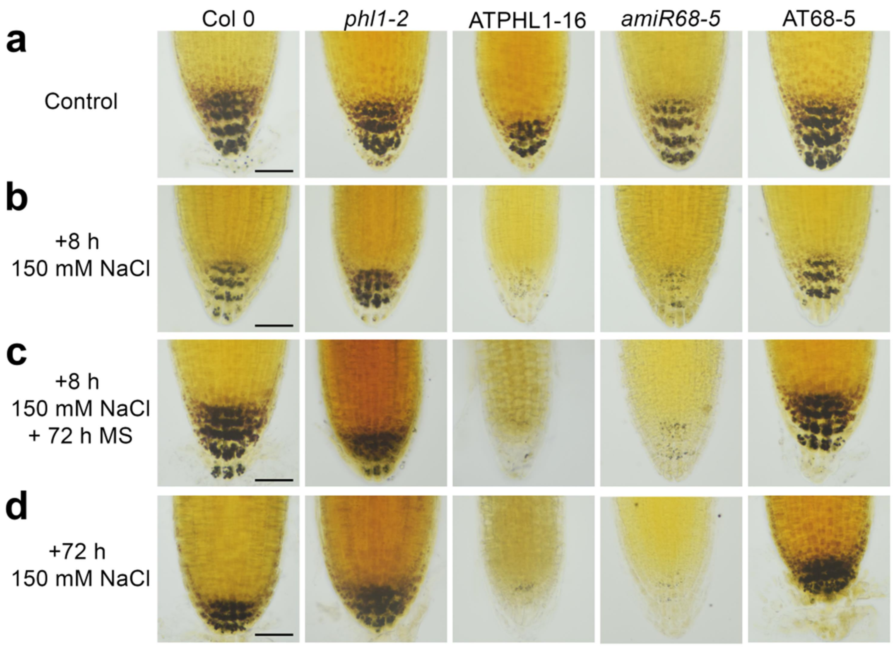


**Figure S7. The adaptation ability to salinity depending on *AtHB23*, *AtPHL1*, and *AtMYB68* levels correlates with the starch granules stage in the root tip**

**a.** Illustrative pictures of root tips (5-day-old) stained with Lugol solution of Col 0, *phl1-2*, OEPHL1-16, *amiR68-5*, and AT68-5 seedlings grown in normal conditions. **b**. The same genotypes after 8 h treatment with 150 mM NaCl. **c**. The roots were transferred to normal conditions or maintained in 150 mM NaCl for additional 72 h (**d**). The black bar represents 50 µm.

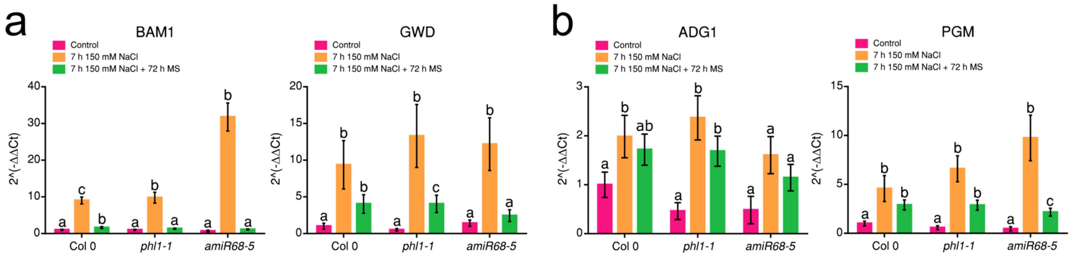


**Figure S8. Starch synthesis and degradation are affected by AtPHL1 and AtMYB68**

Transcript levels of key genes in WT (Col 0), *phl1-1*, and *amiR68–5* plants grown in normal conditions for 5 days treated 7 h with 150 mM NaCl, and placed to recover in MS medium for additional 72 h. Genes assessed participating from degradation were *GWD* and *BAM1* (**a**), and from synthesis *PGM* and *ADG1* (**b**). All the values were normalized with the one obtained in the WT (Col 0). Bars represent SEM. Data were analyzed using a two-way ANOVA considering genotype and treatment. Different letters indicate significant differences (Tukey test, $P < 0.01$).

**Supplementary Table S1**

**Oligonucleotides used for cloning, RT-qPCR or BiFC**

| Oligonucleotide name | Oligonucleotide sequence expressed 5'-3' | Used for construct or analysis of |
|---|---|---|
| I miR68-s | gATTgTTCTAggTTACCAgACggTCTCTCTTTTgTATTCC | 35S:amiR68 |
| II miR68-a | gACCgTCTggTAACCTAgAACAATCAAAgAgAATCAATgA | 35S:amiR68 |
| III miR68*s | gACCATCTggTAACCAAgAACATTCACAggTCgTgATATg | 35S:amiR68 |
| IV miR68*a | gAATgTTCTTggTTACCAgATggTCTACATATATATTCCT | 35S:amiR68 |
| CDSmyb68FBamHI | CCCggATCCATgggAAgAgCACCgTgTTgTgAT | 35S:AtMyb68::GFP |
| CDSmyb68RXbaI | CCCTCTAgACACATgATTTggCgCATTgAAgTA | 35S:AtMyb68::GFP |
| MYB68PROMF | CCCggATCCTgTTCCTTAATCTCAgCTgCCgTgg | PromMYB68::GUS::GFP<br>PromMYB68:MYB68:GUS::GFP |
| MYB68PROMRXhoI | CCCCTCgAgCgTTggCCTTATCACAACACggTgC | PromMYB68::GUS::GFP |
| MYB68RXhoICDS | CCCCTCgAgCACATgATTTggCgCATTgAAgTA | PromMYB68:MYB68:GUS::GFP |
| myb68FBamHI | CCCggATCCATgggAAgAgCACCgTgTTgTgAT | 35S:AtMyb68::GFP |
| myb68RXbaI | CCCTCTAgACACATgATTTggCgCATTgAAgTA | 35S:AtMyb68::GFP |
| AtHB23CDSFw/stop | gggggATCCATgTCTTgTAATAATAATggCTTAgC | 35S:AtHB23:Cherry |
| AtHB23CDSRw/stop | gggCTCgAgTCATCAATTgTATTgTTgCTggTC | 35S:AtHB23:Cherry |
| myb68qPCRf | TTCTTCAATggTTTgggAgCAg | *MYB68* |
| myb68qPCRr | ACTgAggCgTCTCCATAgACTT | *MYB68* |
| ADG1-qPCR-FP | CACCGTCTAAGATGCTTGATGC | *ADG1* |
| ADG1-qPCR-RP | GATGTGCGAGTTTTTCCCAAT | *ADG1* |

| | | |
|---|---|---|
| PGM-qPCR-FP | GTGAAAGAGTATTGGGCGACA | *PGM* |
| PGM-qPCR-RP | CCGTGAACACAAACCGAACA | *PGM* |
| GWD-qPCR-FP | AACGAGAGAGCATACTTCAGC | *GWD* |
| GWD-qPCR-RP | CAATCGGTTTGCTTGGGTAG | *GWD* |
| BAM1-qPCR-FP | ACACGAGCAGATTCTCAAGGC | *BAM1* |
| BAM1-qPCR-RP | TCCCTTCACCCATTTTCTTCA | *BAM1* |
| AtHB23-F | aaaaagcaggctcaATGTCTTGTAATAATAATGGCT | *AtHB23* |
| AtHB23-R | agaaagctgggtaATTGTATTGTTGCTGGTCAAG | *AtHB23* |
| AtMYB68-F | aaaaagcaggctcaATGGGAAGAGCACCGTGTTGT | *MYB68* |
| AtMYB68-R | agaaagctgggtaCACATGATTTGGCGCATTGAA | *MYB68* |
| AtPHL1-F | aaaaagcaggctcaATGACTCTGGCTAATGATTTC | *AtPHL1* |
| AtPHL1-R | agaaagctgggtaATCTTCTCTGACACGTTTCCT | *AtPHL1* |
| HY5-F | aaaaagcaggctcaATGCAGGAACAAGCGACTAGC | *HY5* |
| HY5-R | agaaagctgggtaGACTCGTAAATGTGATAAACCA | *HY5* |
| STO-F | aaaaagcaggctcaATGAAGATACAGTGTGATGTGT | *STO* |
| STO-R | agaaagctgggtaGCCAAGATCAGGGACAATGAA | *STO* |
